# Phosphatidylinositol-4-kinase IIα licenses phagosomes for TLR4 signaling and MHC-II presentation in dendritic cells

**DOI:** 10.1101/2020.01.22.915017

**Authors:** C López-Haber, R Levin-Konigsberg, Y Zhu, J Bi-Karchin, T Balla, S Grinstein, MS Marks, AR Mantegazza

## Abstract

Toll like receptor (TLR) recruitment to phagosomes in dendritic cells (DCs) and downstream TLR signaling are essential to initiate antimicrobial immune responses. However, the mechanisms underlying TLR localization to phagosomes are poorly characterized. We show herein that phosphatidylinositol-4-kinase IIα (PI4KIIα) plays a key role in initiating phagosomal TLR4 responses in murine DCs by generating a phosphatidylinositol-4-phosphate (PtdIns4P) platform conducive to the binding of the TLR sorting adaptor TIRAP. PI4KIIα is recruited to LPS-containing phagosomes in an adaptor protein AP-3 dependent manner, and both PI4KIIα and PtdIns4P are also detected on phagosomal membrane tubules. Knockdown of PI4KIIα – but not of the related PI4KIIβ – impairs TIRAP and TLR4 localization to phagosomes, reduces proinflammatory cytokine secretion, and impairs phagosomal tubule formation and MHC-II presentation. Phagosomal TLR responses in PI4KIIα-deficient DCs are restored by re-expression of wild-type PI4KIIα, but not of variants lacking kinase activity or AP-3 binding. Our data indicate that PI4KIIα is an essential regulator of phagosomal TLR signaling in DCs by ensuring optimal TIRAP recruitment to phagosomes.

## INTRODUCTION

Signaling by pattern recognition receptors such as Toll-like receptors (TLRs) is essential to initiate immune responses (Medzhitov & Janeway, 2002). In addition, compartmentalization of TLR signaling is a fundamental mechanism that allows discrimination between a myriad of self and foreign stimuli that may pose different levels of threat (Barton & Kagan, 2009). One of the cellular compartments where TLR signaling is particularly important is the phagosome, a lysosome-related organelle formed in phagocytic cells such as dendritic cells (DCs) upon the capture of a particulate target, such as a bacterium (Flannagan et al, 2012; Mantegazza & Marks, 2016). Particularly in DCs, which serve as the main interface between innate and adaptive immunity, the phagosome becomes an autonomous TLR-sensing and -signaling platform that contains all the machinery required to process the captured material, load resulting peptides into MHC molecules and form phagosomal tubules (phagotubules) that favor antigen presentation to T cells (Hoffmann et al, 2012; Mantegazza et al, 2014; Underhill & Goodridge, 2012), and extends proinflammatory responses initiated by plasma membrane TLRs by the acquisition of an additional pool of intracellular TLRs. This second wave of TLR signaling from the phagosome is focused on a single potentially harmful particulate entity. It is therefore essential to keep phagosomal identity different from the plasma membrane, where sensing reflects a broader spectrum of stimuli (Blander & Sander, 2012), and also from endosomes, which can bear soluble cargo that may be less harmful. In addition, preserving phagosomal identity and autonomy is essential in DCs for the initiation of adaptive immune responses.

While TLR signaling pathways have been extensively studied, less is known about how TLR recruitment to phagosomes in DCs is regulated to optimize the immune response. Our current understanding of TLR recruitment to phagosomes in DCs was largely shaped by our analyses of a mouse model of Hermansky-Pudlak syndrome type 2 (HPS2) (Mantegazza et al, 2012). HPS2 is caused by inactivating mutations in the β3A subunit of AP-3, an adaptor protein that binds to cytoplasmic signals in transmembrane proteins and mediates their trafficking from early endosomes to lysosomes or lysosome-related organelles (Dell’Angelica et al, 1999). HPS2 is characterized by immunodeficiency among other symptoms (Bowman et al, 2019; Dell’Angelica & Bonifacino, 2019; Mantegazza & Marks, 2016). While the role of AP-3 in T cell and plasmacytoid DC function (Blasius et al, 2010; Clark et al, 2003; Sasai et al, 2010) may explain recurrent viral infections in HPS2, defects in lipid antigen presentation and granulopoiesis (Briken et al, 2002; Fontana et al, 2006; Sugita et al, 2002), together with our observations that AP-3 is required for optimal secretion of proinflammatory cytokines and MHC-II presentation of phagocytosed antigen to T cells (Mantegazza et al, 2012), may explain defective anti-bacterial immunity in HPS2. The impaired immune responses in AP-3 deficient conventional DCs are at least in part due to the reduced recruitment of TLR4 and other TLRs to maturing phagosomes from an endosomal pool (Mantegazza et al, 2012). However, it is not known whether AP-3 supports TLR4 localization to phagosomes by direct binding and incorporation into phagosome-bound vesicles or by an indirect mechanism.

We and others showed that TLR4 signaling to a set of downstream immune responses in DCs requires the engagement of the TLR signaling adaptor MyD88 (Blander & Medzhitov, 2004; Mantegazza et al, 2014). MyD88 binding to TLR4 is facilitated by the TLR sorting adaptor Toll-IL1 receptor (TIR) domain-containing adaptor protein (TIRAP; aka MAL) (O’Neill & Bowie, 2007), which also binds TLR4 and allows the formation of a multimolecular signaling platform known as the MyDdosome (Gay et al, 2011). Therefore, TLR4 requires TIRAP to dictate the cellular location where MyD88-dependent signaling will start. Importantly, TIRAP binding to cellular membranes depends on the presence of polyphosphorylated inositide-(or phosphoinositide-) enriched domains, highlighting the importance of phosphoinositides as essential mediators of TLR4 positioning at sites of future signal transduction (Bonham et al, 2014; Kagan & Medzhitov, 2006).

Phosphoinositides play an essential role in cellular membrane physiology by regulating membrane curvature, protein recruitment and vesicular trafficking and therefore defining membrane identity (Hammond et al, 2012; Janmey et al, 2018; Krauss & Haucke, 2007; Tan & Brill, 2014). The type II lipid kinases phosphatidylinositol-4-kinase IIα (PI4KIIα) and IIβ (PI4KIIβ), (Craige et al, 2008; Salazar et al, 2005)are distinctly encoded enzymes that phosphorylate phosphatidylinositol (PtdIns) at position 4, generating PtdIns4P at the plasma membrane, Golgi, trans Golgi network (TGN) and endolysosomes (Minogue, 2018). In particular, PI4KIIα is abundant on membranes of the endolysosomal network (Balla & Balla, 2006; Balla et al, 2002) notably in a vesicular pool rich in AP-3 (Salazar et al, 2005). Importantly, PI4KIIα binds AP-3 directly via a classical dileucine-based sorting signal and is thereby delivered to lysosomes and lysosome-related organelles (Craige et al, 2008). In macrophages, PI4KIIα and its product PtdIns4P are also required for phagosome maturation and resolution (Jeschke et al, 2015; Levin et al, 2017; Levin-Konigsberg et al, 2019). However, a possible link between PI4KIIα and phagosomal TLR signaling has not been investigated.

Considering the connection between PI4KIIα and AP-3, and the requirement for phosphoinositides in TIRAP/MyD88-dependent TLR4 signaling, we investigated whether phagosomal TLR4 signaling in mouse DCs is dependent on the lipid kinase PI4KIIα and its product PtdIns4P. We show herein that PI4KIIα, but not the related PI4KIIβ, is required for the production of PtdIns4P on phagosomes in DCs, for optimal TIRAP binding to phagosomes, and for downstream TLR4 responses. PI4KIIα recruitment to phagosomes is in turn dependent on AP-3 function, which may therefore at least partly explain the defective anti-bacterial responses in AP-3 deficient mouse DCs and HPS2 patients. Finally, we show that, compared to macrophages, PI4KIIα plays a differential role in DC phagosomes that reflects DC specialization in antigen presentation and ensures the preservation of phagosome identity and autonomous signaling.

## RESULTS

### 1. PI4KIIα is recruited to DC phagosomes in an AP-3-dependent manner

AP-3 recognizes accessible cytosolic targeting sequences on transmembrane proteins and mediates their trafficking to lysosomes or lysosome-related organelles. The consensus sequences YXXϕ or [DE]XXXL[LI] are recognized by the µ3 and σ3-δ subunits pf AP-3, respectively (Bonifacino & Traub, 2003; Owen & Luzio, 2000). TLR4 bears a YDAF sequence in the cytoplasmic TIR domain. However, the TLR4 crystal structure suggests that the tyrosine is not accessible for intermolecular interactions (Wang et al, 2016). Consistent with this, we could not detect direct binding of the TLR cytoplasmic domain to the AP-3 µ3 subunit by yeast-two hybrid analysis (**Figure EV1**) under the same conditions that allowed binding of the cytoplasmic tail of human TGN38 to the µ subunits of AP-1, −2 and −3, as expected (Ohno et al, 1998; Ohno et al, 1995) (**Figure EV1**). Although these data do not exclude a direct interaction through other subunits, the absence of the most likely µ3 - TLR4 interaction led us to investigate whether additional effectors might favor AP-3-dependent TLR4 recruitment to phagosomes in DCs.

TLR4 signaling complexes are found on lipid microdomains enriched in phosphoinositides. Because PtdIns4P and PI4KIIα are present on maturing phagosomes in macrophages (Jeschke et al, 2015; Levin et al, 2017), and AP-3 directly interacts with PI4KIIα in neurons and other cell types (Craige et al, 2008; Salazar et al, 2005), we decided to investigate the role of PI4KIIα and its product PtdIns4P in TLR4 recruitment to DC phagosomes. We first investigated if PI4KIIα and PtdIns4P were present on phagosomes in DCs and whether PI4KIIα recruitment was dependent on AP-3. Bone marrow-derived DCs from wild-type (WT) and Ap3b^pe/pe^ mice that lack AP-3 in non-neuronal cell types (AP-3^-/-^) were pulsed with magnetic beads coated with the TLR4 ligand LPS and chased over time after phagocytosis. Phagosomes were then isolated and analyzed by immunoblotting (Figures 1A, B). Phagosomes from WT DCs showed an increased acquisition of endogenous PI4KIIα over 2 h. In contrast, whereas cellular expression of PI4KIIα in WT and AP-3^-/-^ DCs was similar, PI4KIIα recruitment to phagosomes in AP-3^-/-^ DCs was significantly reduced (Figures 1A, B). To confirm these results, we expressed PI4KIIα-GFP in DCs by recombinant retroviral transduction of bone marrow (transduction efficiency was similar in WT and AP-3^-/-^ DCs; **Figure EV2A**), pulsed differentiated DCs with LPS-coated polystyrene beads, and tested for PI4KIIα-GFP recruitment to DC phagosomes by flow cytometry of isolated phagosomes (Figures 1C, D and **EV2B**) or by live cell imaging (Figure 1E-G). Like endogenous PI4KIIα, PI4KIIα-GFP was increasingly recruited to phagosomes over 2 h after phagocytosis in WT DCs, while recruitment to AP-3^-/-^ DCs was significantly impaired (Figure 1C, D); after normalizing to the percent of PI4KIIα-GFP positive DCs (**Figure EV2A**), PI4KIIα-GFP was recruited to 90% of phagosomes in WT DCs but only 20% in AP-3^-/-^ DCs at 120 min (Figure 1D). Consistent with these results, PI4KIIα-GFP recruitment to phagosomes containing polystyrene beads coated with LPS and Texas red-conjugated ovalbumin (LPS/OVA-TxR) was significantly impaired in AP-3^-/-^ DCs, as observed by live cell imaging on transduced DCs (97±3% in WT DCs vs. 12±3% in AP-3^-/-^ after 120 min; Figure 1E-G**, Movies EV1, 2**). Together, these data indicate that AP-3 is required for optimal recruitment of PI4KIIα to phagosomes. Noteworthy, PI4KIIα-GFP was also recruited to phagotubules in maturing phagosomes from WT DCs (Figure 1E, time 120 min; **Movies EV3, 4**).

**Figure 1.**
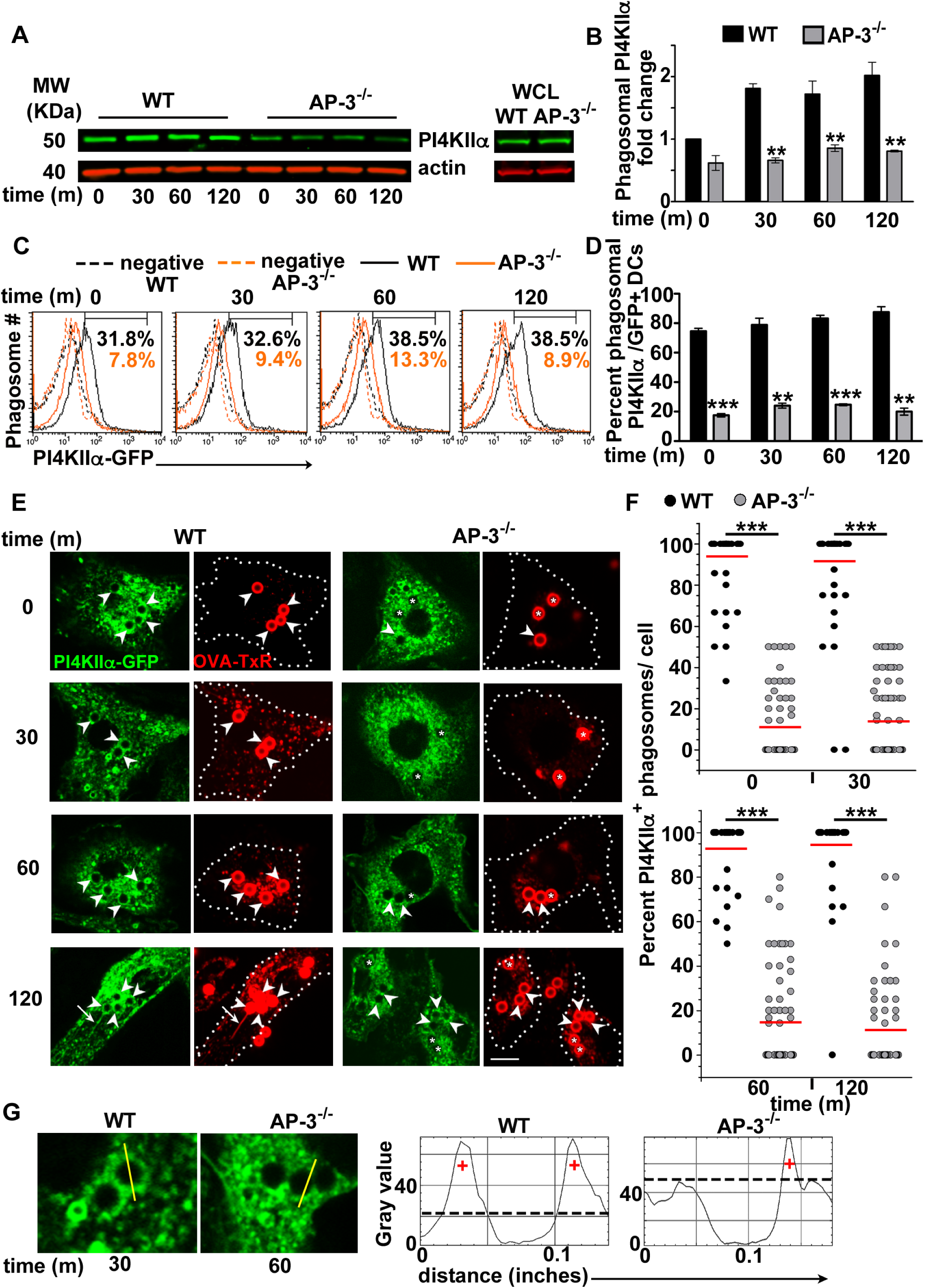
PI4KIIα is recruited to DC phagosomes in an AP-3-dependent manner. WT and AP-3^-/-^ BMDCs that had been non-transduced (A, B) or transduced with retroviruses encoding PI4KIIα-GFP (C-G), were pulsed with LPS-coated magnetic beads (A, B), LPS-coated polystyrene beads (C, D) or LPS/OVA-TxR-coated polystyrene beads (E-G) and chased as indicated. A. Purified phagosomes (*left*) or whole cell lysates (*right*) were analyzed by SDS-PAGE and immunoblotting for PI4KIIα and actin. Shown are the relevant bands for each blot. B. Quantification of band intensities for phagosomal PI4KIIα from three independent experiments, showing fold change relative to WT time 0 and normalized to actin (mean ± SD). (C, D) Phagosomes from non-transduced or PI4KIIα-GFP-expressing WT and AP-3^-/-^ BMDCs were purified and analyzed by flow cytometry. GFP-positive BMDCs were not previously sorted. C. Shown are histogram plots of a representative experiment with the percentages of gated phagosomes that were GFP positive (signal over phagosomes from non-transduced cells) indicated. Solid black lines, WT; solid orange lines, AP-3^-/-^; dashed lines, non-transduced controls. (D) Data (mean ± SD) from three independent experiments performed in duplicate were normalized to the percent of transduced BMDCs (see **Supp. Fig. 2A**). (E-G) WT and AP-3^-/-^ BMDCs expressing PI4KIIα-GFP were analyzed by live cell imaging. E. Representative images at indicated times after phagocytosis. Dotted white lines, cell outlines; arrowheads, PI4KIIα-GFP positive phagosomes; asterisks, PI4KIIα-GFP negative phagosomes; arrows, phagotubules. Scale bar: 9 µm. F. Data from three independent experiments, 20 cells per experiment, are presented as percent of PI4KIIα-GFP**^+^** phagosomes per cell. Black dots, WT; grey dots, AP-3^-/-^; solid red lines, means. G. Quantification was performed as shown on representative images, drawing a line across the phagosomes (*left*) and analyzing the line plot with ImageJ (*right;* +, positive signal). Phagosomes containing GFP positive puncta, even partially in the total phagosomal membrane surface (mostly in the case of AP-3^-/-^ BMDCs) were considered positive. **p<0.01; ***p<0.001.

### 2. PI4KIIα accumulates PtdIns4P on early and maturing phagosomes

To test whether PtdIns4P was produced on DC phagosomes concomitantly with PI4KIIα recruitment, we expressed GFP-tagged P4M-SidMx2 (P4Mx2) – a probe for PtdIns4P derived from the SidM domain of *Legionella pneumophila* (Hammond et al, 2014) – in DCs and analyzed cells over time after phagocytosis of LPS/OVA-TxR beads using live imaging. In addition, we assessed the contribution of PI4KIIα and the related PI4KIIβ – which does not bind AP-3 – to the production of PtdIns4P on phagosomes by transducing bone marrow precursors with shRNA targeted to each PI4KII isoform or a non-target control shRNA (**Figure EV3A,** Figure 2A). Knockdown was specific for each PI4KII isoform. PI4KIIα shRNA #2 was more effective than #1 and was used in subsequent experiments. Retroviral transduction did not significantly affect DC differentiation (**Figure EV3B**), and phagocytic capacity was not affected by the protein knockdowns (**Figure EV3C**). DC maturation in response to LPS was consistently reduced by shRNA transduction but comparable between the different shRNA treatments (**Figure EV3D**). The GFP-P4Mx2 probe detected PtdIns4P on the plasma membrane and in an intracellular pool (as described in other cells (Hammond et al, 2014)) (Figure 2A). In addition, after engulfment of LPS/OVA-TxR beads, PtdIns4P was found on nascent phagosomes in all DC types. In addition, PtdIns4P was present in early and maturing phagosomes and in phagotubules in shRNA control DCs (Figure 2A). However, DCs knocked down for PI4KIIα showed significantly reduced accumulation of PtdIns4P on early and maturing phagosomes (98% in control DCs versus 0% in PI4KIIα knockdown DCs between 30 and 120 min; Figure 2B), while the plasma membrane signal was not affected (Figure 2A). Conversely, DCs knocked down for PI4KIIβ showed reduced binding of GFP-P4Mx2 to the plasma membrane, while accumulation of PtdIns4P on phagosomes (99±1% between 30 and 120 min) and phagotubules was unaffected (Figure 2A). Moreover, and consistent with the reduced recruitment of PI4KIIα to phagosomes in AP-3^-/-^ DCs (Figure 1), PtdIns4P formation was significantly reduced in AP-3-deficient DCs (2±2% between 30 and 120 min, Figure 2B). These data suggest that AP-3 and its cargo protein PI4KIIα are required for PtdIns4P formation on phagosomes, while PI4KIIβ may primarily contribute to the plasma membrane PtdIns4P pool in DCs.

**Figure 2.**
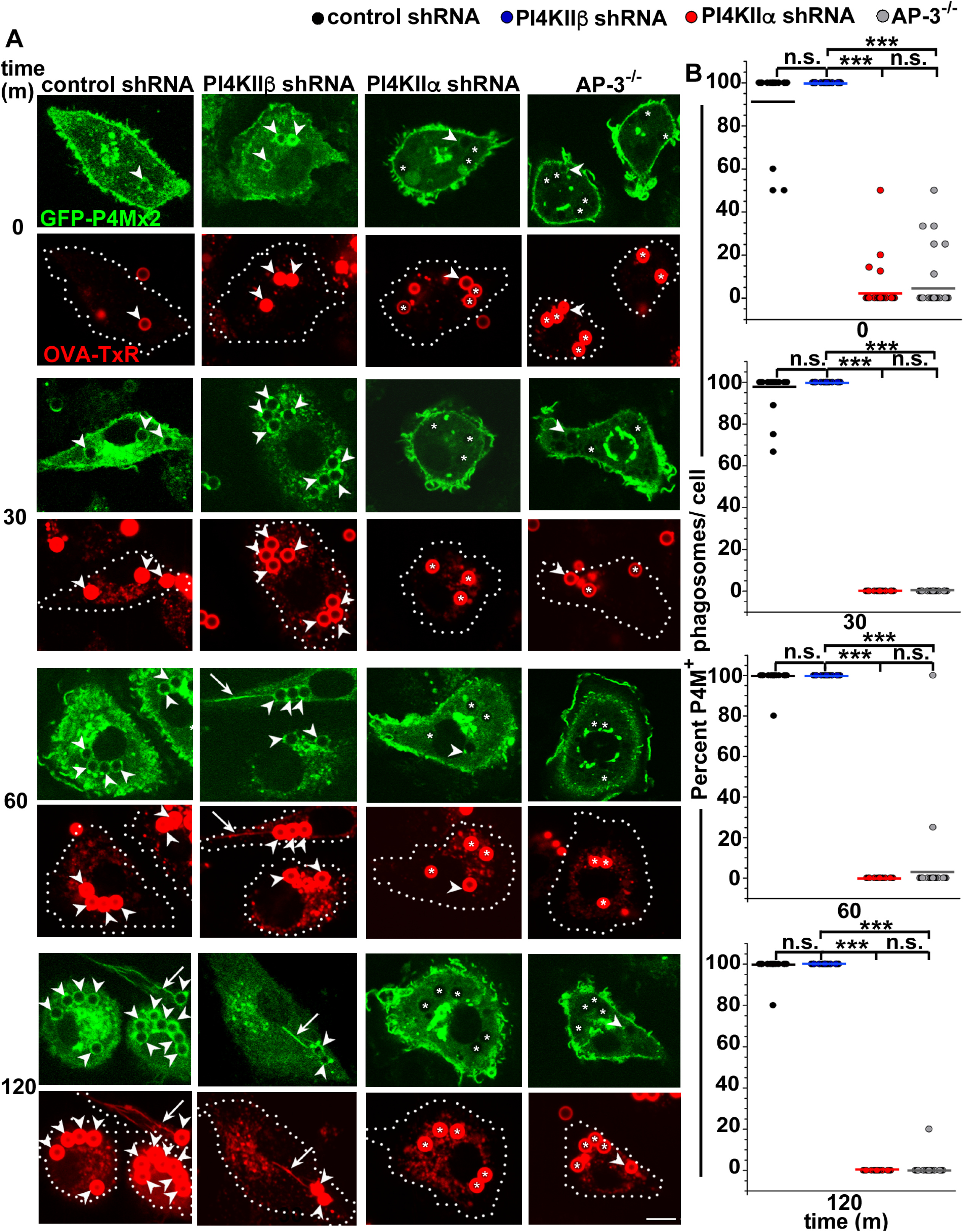
PI4KIIα is needed to accumulate PtdIns4P on early and maturing phagosomes. WT BMDCs were transduced with retroviruses encoding GFP-P4Mx2 and with lentiviruses encoding non-target (control), PI4KIIβ or PI4KIIα shRNAs, and AP-3^-/-^ BMDCs were transduced only with retroviruses encoding GFP-P4Mx2. DCs were pulsed with LPS/OVA-TxR-coated beads, chased as indicated and analyzed by live cell imaging. A. Representative images. Dotted white lines, cell outlines; arrowheads, GFP-P4Mx2 positive phagosomes; asterisks, GFP-P4Mx2 negative phagosomes; arrows, phagotubules. B. Data from three independent experiments, 20 cells per experiment, are presented as percent of GFP-P4Mx2**^+^** phagosomes per cell. Black dots, non-target control shRNA; blue dots, PI4KIIβ shRNA; red dots, PI4KIIα shRNA; grey dots, AP-3^-/-^; solid color lines, means. Scale bar: 9 µm. ***p<0.001; n.s., not significant.

### 3. PI4KIIα is required for optimal phagosomal TLR4 signaling and MHC-II presentation of phagocytosed antigen in DCs

We and others showed that TLR4 activation via its signaling adaptor MyD88 on phagosomes triggers a signaling cascade that leads to proinflammatory cytokine production, phagotubule formation and MHC-II presentation of phagocytosed cargo in DCs (Blander & Medzhitov, 2006; Mantegazza et al, 2012; Mantegazza et al, 2014). To test whether these responses require PI4KIIα, we probed for TLR4/MyD88 responses in DCs derived from bone marrow transduced with non-target shRNA or shRNA to PI4KIIα, PI4KIIβ or the SNARE protein Sec22b (Cebrian et al, 2011) as an additional control (**Figure EV3A**). We first analyzed the production of the proinflammatory IL-6 by DCs pulsed with LPS-coated beads (relative to uncoated beads as a negative control) by ELISA analysis of cell supernatants. IL-6 levels were significantly reduced in supernatants of PI4KIIα knockdown DCs compared to those from PI4KIIβ knockdown DCs or DCs transduced with control shRNAs (Figure 3A). To test for phagotubule formation, we analyzed DCs by live cell imaging 2.5 h after exposure to LPS/OVA-TxR beads. Whereas PI4KIIβ knockdown DCs or DCs treated with control shRNAs exhibited robust phagotubule formation, PI4KIIα knockdown severely impaired the formation of phagotubules, as also observed in AP-3^-/-^ DCs (Figure 3B, C**, Movies EV5-7**; see also Figures 1E and 2A, time 120 min). These data show that both TLR-induced proinflammatory cytokine secretion and phagotubule formation require PI4KIIα.

**Figure 3.**
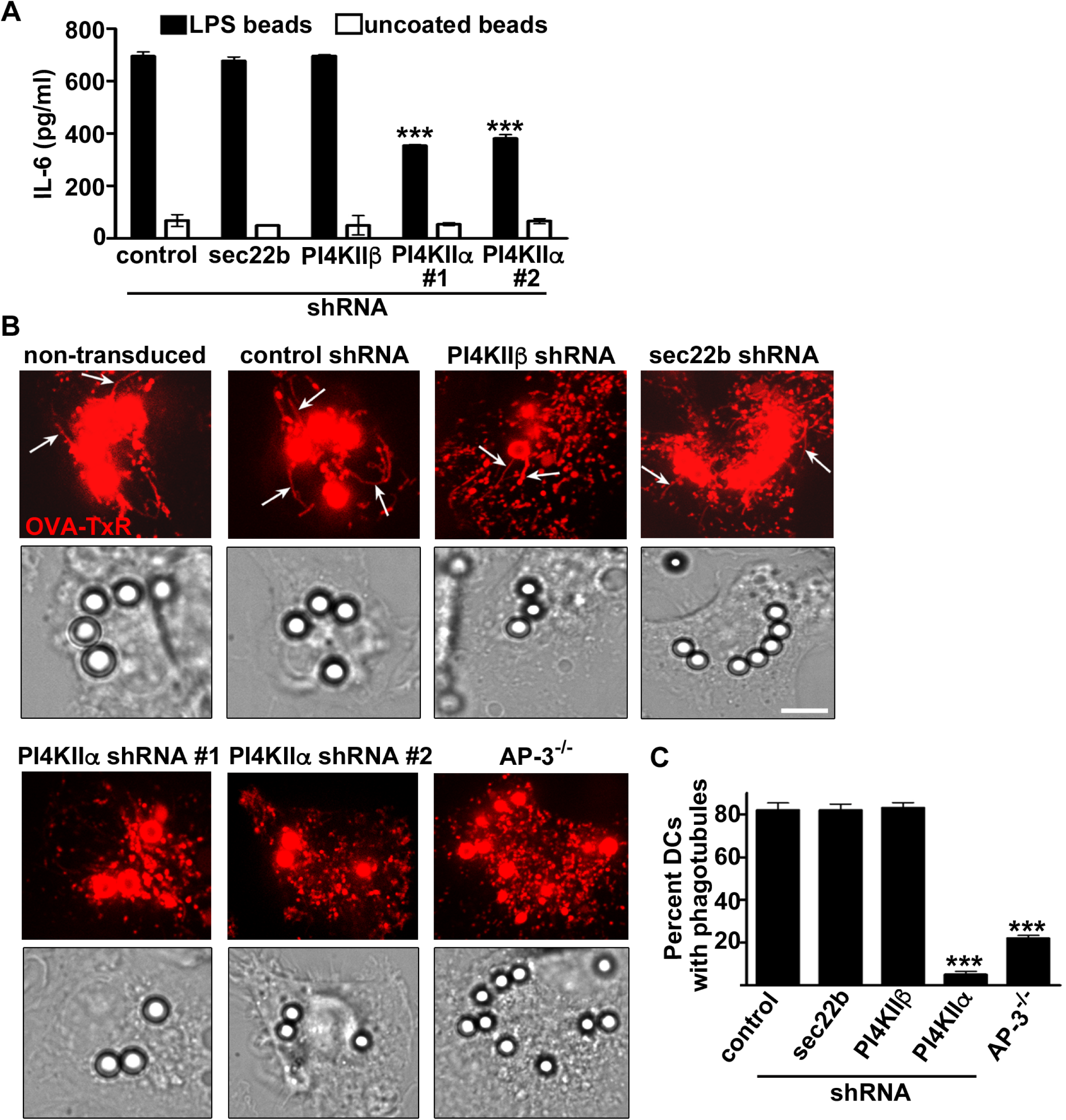
PI4KIIα is required for optimal phagosomal TLR4 signaling. WT BMDCs were non-transduced (-) or transduced with lentiviruses encoding non-target (control), PI4KIIβ, Sec22b or either of two PI4KIIα shRNAs, and AP-3^-/-^ BMDCs were untransduced. DCs were pulsed with uncoated or LPS-coated polystyrene beads (A) or LPS/OVA-TxR-coated beads (B, C). A. IL-6 released into the supernatants was measured by ELISA after a 3 h chase. Results represent mean ± SD of three experiments each performed in triplicate. (B, C). BMDCs were analyzed by live cell imaging after a 2.5 h chase. B. Arrows indicate phagotubules. Scale bar: 9 µm. C. The percentage of BMDCs presenting phagotubules (tubules ≥1 µm emerging from phagosomes) in three independent experiments, 20 cells per experiment, is shown. ***p<0.001. No significance is not indicated.

Finally, we assessed MHC-II presentation of phagocytosed antigen using beads coated with the Eα_52-68_ peptide (subsequent presentation of this peptide by the MHC-II molecule I-A^b^ at the cell surface can be detected by the YAe antibody (Rudensky et al, 1991)). DCs were pulsed with EαGFP-coated beads or with soluble EαGFP (to monitor presentation of soluble antigen) or Eα_52-68_ peptide alone (a control that binds to surface I-A^b^); all preparations contained LPS to stimulate TLR4. As expected from the reduced DC maturation observed in shRNA-transduced cells (**Figure EV3D**), MHC-II expression on transduced DCs was reduced compared to non-transduced DCs but similar between the shRNA treatments (except for Sec22b shRNA, which was reduced more and thus not included in these assays; **Figure EV4**). As observed for AP-3^-/-^ DCs (Mantegazza et al 2012), formation of cell surface Eα:I-A^b^ complexes 6 h following exposure to Eα_52-68_ peptide or soluble EαGFP was not significantly affected by PI4KIIα shRNA compared to the other shRNAs (Figure 4A, B). By contrast, surface presentation of peptide:MHC-II complexes 6 h after phagocytosis of EαGFP-coated beads was significantly reduced by expression of PI4KIIα shRNA, but not PI4KIIβ or non-target control shRNAs (Figure 4C, D). To test if MHC-II presentation correlated with CD4+ T cell responses, DCs pulsed with the ovalbumin (OVA)-derived OVA_323-329_ peptide, soluble OVA/LPS or OVA/LPS-coated beads were co-cultured with OVA_323-329_-specific, I-A^b^-restricted OT-II T cells. OT-II cell activation, measured by CD69 expression and IL-2 production, by DCs pulsed with peptide alone or soluble OVA was similar whether DCs expressed PI4KIIα, PI4KIIβ or control shRNAs (Figure 4E-H). In contrast, OT-II cell activation by DCs pulsed with OVA/LPS beads was significantly reduced in PI4KIIα shRNA-expressing DCs relative to cells expressing other shRNAs (Figure 4I, J). These data indicate that PI4KIIα promotes MHC-II presentation of antigen following phagocytosis but not endocytosis. This is in agreement with our observations that PI4KIIα is required for PtdIns4P formation on phagosomes, but not on the plasma membrane (Figure 2).

**Figure 4.**
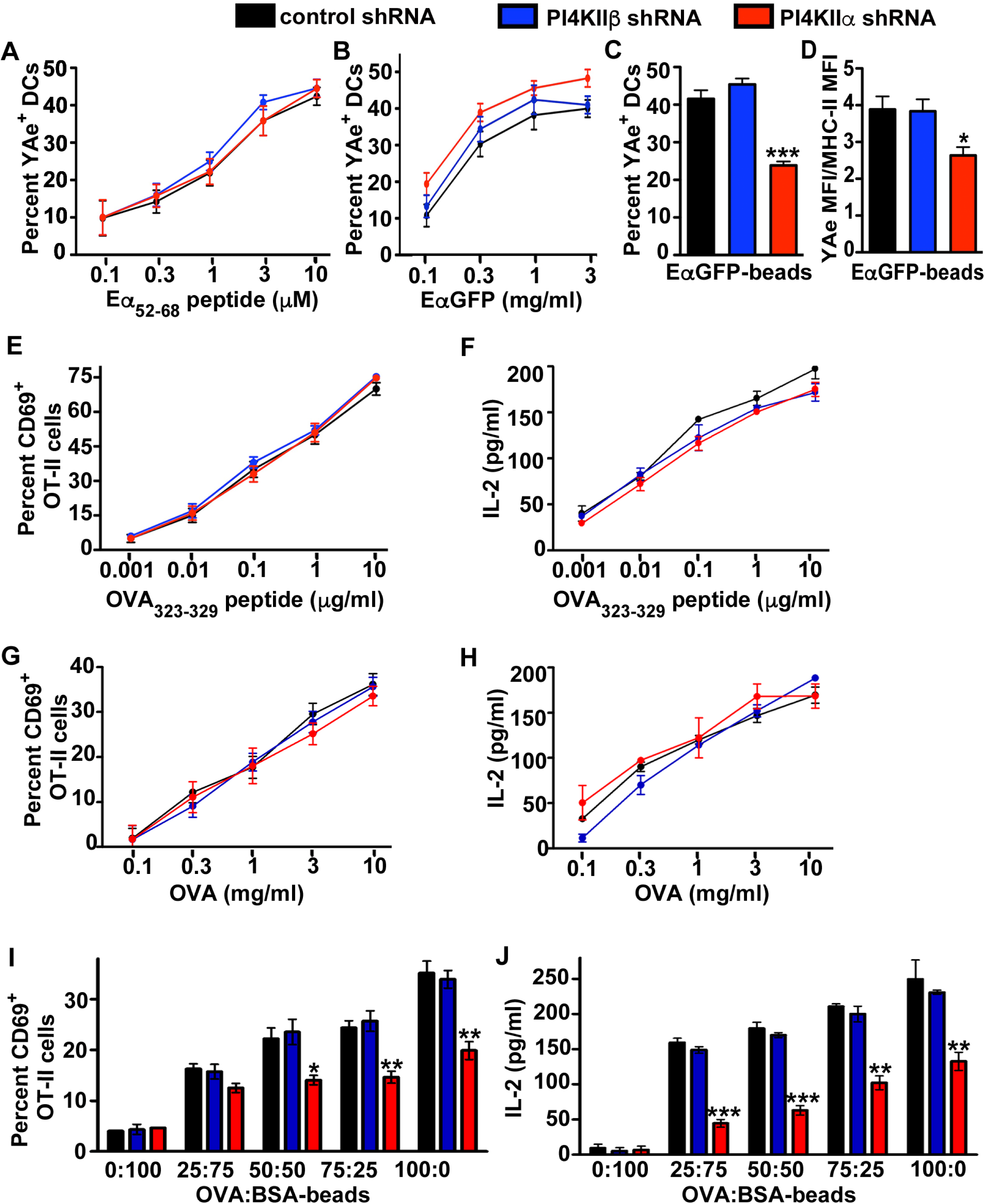
PI4KIIα is required for optimal MHC-II presentation of phagocytosed antigen. WT BMDCs transduced with lentiviruses encoding non-target (control), PI4KIIβ or PI4KIIα shRNAs were pulsed with Eα_52-68_ peptide (A), soluble EαGFP fusion protein (B), EαGFP-coated beads (C, D), OVA_323-329_ peptide (E, F), soluble OVA (G, H) or OVA:BSA-coated beads (I, J) and chased for 6h. (A-D). Surface expression of Eα_52-68_:I-A^b^ complexes was analyzed by flow cytometry using YAe antibody. (A-C). Shown are the percentages of CD11c**^+^** BMDCs that were Eα_52-68_:I-A^b**+**^. D. Shown is the YAe MFI normalized to MHC-II MFI (see **Supp. Figure S3D**). (E-J). Pulsed BMDCs were fixed and cocultured with pre-activated OT-II cells for 18h. (E, G, I). CD69 expression by OT-II cells was assessed by flow cytometry. Shown are the percentages of vb5**^+^** T cells that expressed CD69. (F, H, J). IL-2 production by OT-II cells was measured by ELISA on the coculture supernatants. Data in all panels represent mean ± SD of three experiments performed in duplicate. *p<0.05; **p<0.01; ***p<0.001. No significance is not indicated.

### 4. PI4KIIα phagosomal function requires the AP-3-sorting motif and kinase activity

In order to assess if PI4KIIα recruitment and consequent phagosomal TLR signaling required AP-3 binding and kinase activity, we tested whether PI4KIIα mutants could rescue these phenotypes in PI4KIIα knockdown cells. Bone marrow cells were transduced with retroviruses encoding either human WT PI4KIIα-GFP, the AP-3 sorting mutant (L61,62A – which does not bind AP-3 –) or the kinase-inactive mutant (D308A), all previously characterized (Craige et al, 2008), and subsequently transduced with lentiviruses encoding murine PI4KIIα shRNA; control cells were transduced only with non-coding shRNA lentiviruses (Figure 5A). Transduced DCs were then pulsed with LPS/OVA-TxR beads and analyzed by live cell imaging after 2.5 h. Phagosomal recruitment of both the AP-3 sorting mutant and the kinase-inactive mutant was significantly impaired compared to WT PI4KIIα-GFP (WT, 98±1%; D308A, 24±3%; L61,62A, 10±2%; Figure 5B, C), consistent with previous reports for PI4KIIα-GFP recruitment to lysosomes and lysosome related organelles in other cells (Craige et al, 2008; Salazar et al, 2005). Moreover, only WT PI4KIIα-GFP restored phagotubule formation in knockdown DCs and was itself recruited to phagosomal tubules (Figure 5B). WT PI4KIIα-GFP but not the kinase inactive or AP-3 binding mutants also restored proinflammatory cytokine production (Figure 5D) and the ability to present phagocytosed Eα on MHC-II molecules (Figure 5E). Thus, both AP-3 binding and kinase activity are required for PI4KIIα recruitment to phagosomes and consequent phagosomal function in TLR4 signaling.

**Figure 5.**
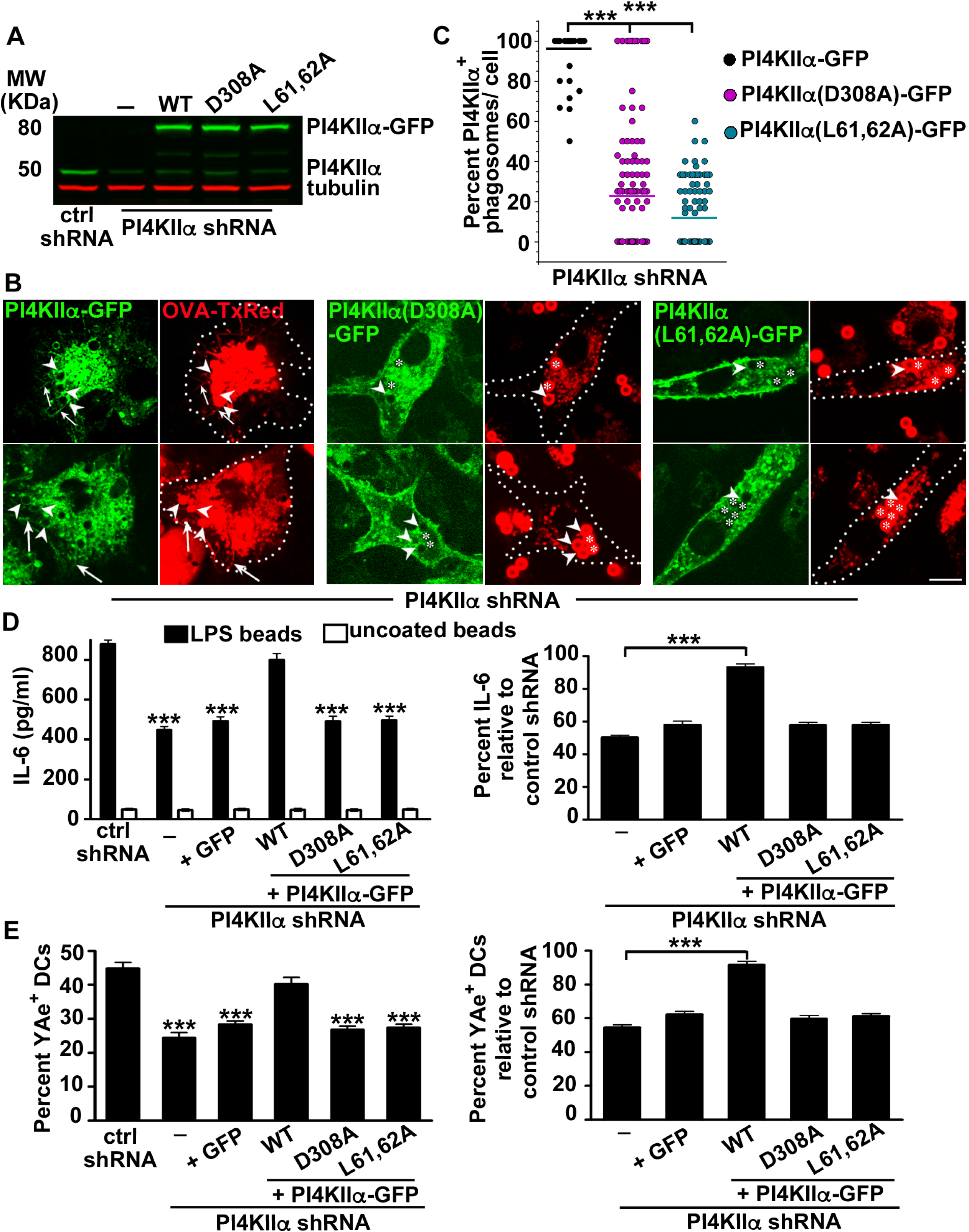
PI4KIIα function on phagosomes requires the AP-3 sorting motif and kinase activity. WT BMDCs that were non-transduced (-) or transduced first with retroviruses encoding GFP, human WT PI4KIIα-GFP, PI4KIIα(D308A)-GFP or PI4KIIα(L61,62A)-GFP, were then transduced with lentiviruses encoding mouse non-target (ctrl) or PI4KIIα shRNAs. Cells were left untreated (A) or pulsed with LPS/OVA-TxR-coated beads (B, C), uncoated or LPS-coated polystyrene beads (D), or EαGFP-coated beads (E). A. Whole cell lysates were analyzed by SDS-PAGE and immunoblotting for PI4KIIα, GFP and tubulin. Shown are the relevant bands for each blot. (B, C). BMDCs were analyzed by live cell imaging after a 2.5 h chase. B. Representative images. Dotted white lines, cell outlines; arrowheads, PI4KIIα-GFP positive phagosomes; asterisks, PI4KIIα-GFP negative phagosomes; arrows, phagotubules. C. Data from three independent experiments, 20 cells per experiment, are presented as percent of PI4KIIα-GFP**^+^** phagosomes per cell. Black dots, WT; purple dots, PI4KIIα(D308A)-GFP; tidal dots, PI4KIIα(L61,62A)-GFP; solid color lines, means. Scale bar: 9 µm. D. IL-6 released into the supernatants after 3 h was measured by ELISA. *Left panel*, representative experiment performed in triplicates. *Right panel*, IL-6 values from three independent experiments performed in triplicate are shown as percent of values for BMDCs treated with non-target (ctrl) shRNA, as a representation of phenotypic rescue (mean ± SD). E. Surface expression of Eα_52-68_:I-A^b^ complexes was analyzed by flow cytometry using YAe antibody. *Left panel*, shown are the percentages of CD11c**^+^** BMDCs that were Eα_52-68_:I-A^b**+**^ in a representative experiment. *Right panel*, YAe values from two independent experiments performed in duplicate are shown as percent of values for BMDCs treated with non-target (ctrl) shRNA, as a representation of phenotypic rescue (mean ± SD). (D, E). Significance relative to non-target shRNA-treated WT control (*left panels*) or PI4KIIα shRNA-treated DCs (-) (*right panels*) is indicated. ***p<0.001. No significance is not indicated.

### 5. PI4KIIα promotes TLR4 accumulation on phagosomes in DCs

To test whether TLR4 localization to phagosomes required PI4KIIα, we pulsed DCs with LPS-coated beads and analyzed TLR4 presence on isolated phagosomes by immunoblotting and flow cytometry using two different antibodies. Immunoblotting showed that cellular TLR4 expression was similar in DCs treated with control, PI4KIIα or PI4KIIβ shRNAs (Figure 6A, *left panel*). In both assays, TLR4 was increasingly accumulated on phagosomes over time after phagocytosis in control and PI4KIIβ knockdown DCs, as we had shown before (Mantegazza et al, 2012), but not in cells knocked down for PI4KIIα (Figure 6A-D). This was true whether data were analyzed for total TLR4 content on phagosomes by immunoblotting (Figure 6A, C) or by percentage of phagosomes harboring TLR4 (Figure 6B, D). These data indicate that TLR4 localization to phagosomes requires PI4KIIα.

**Figure 6.**
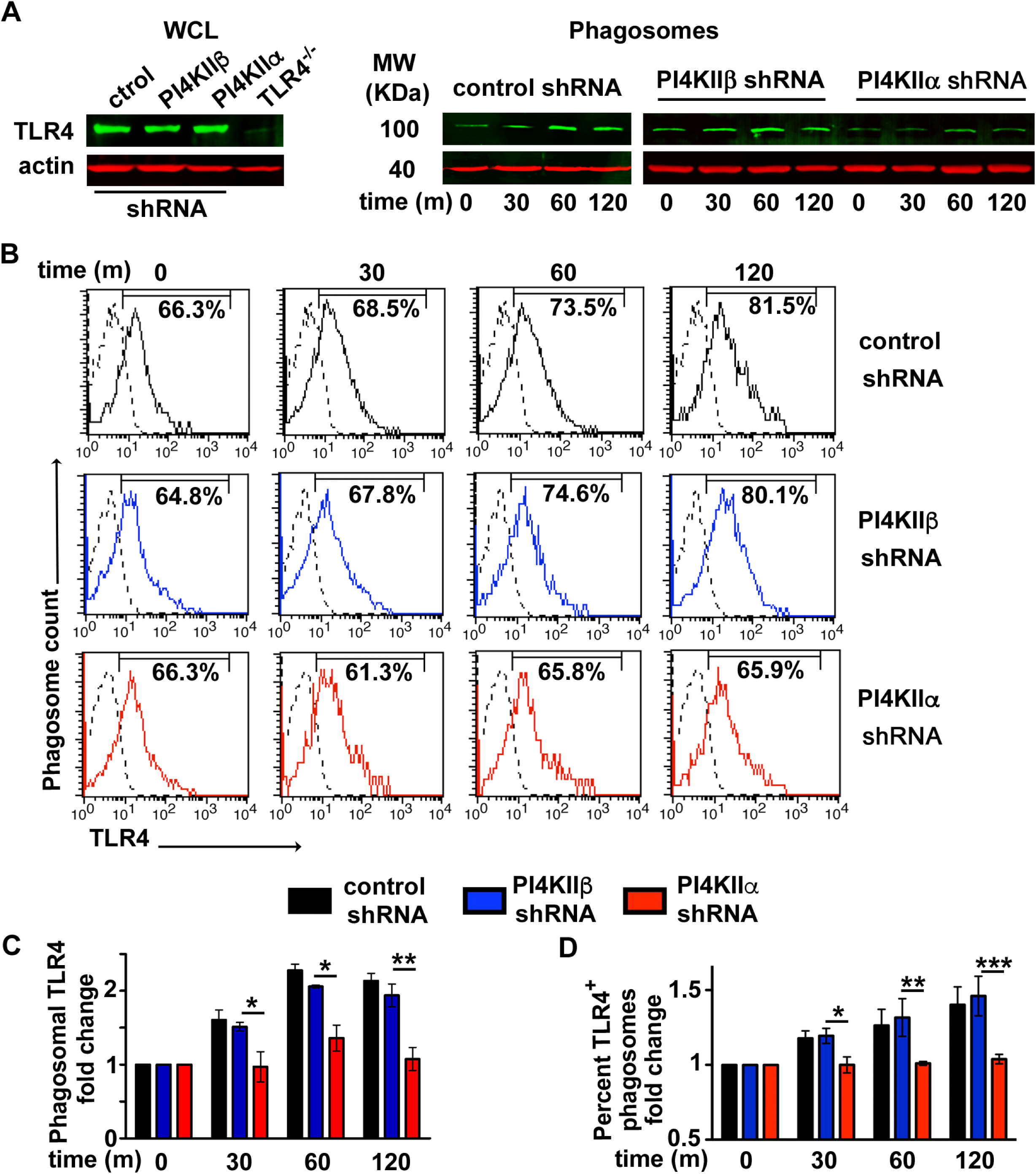
PI4KIIα promotes TLR4 recruitment to phagosomes in DCs. WT BMDCs transduced with lentiviruses encoding non-target (control), PI4KIIβ or PI4KIIα shRNAs were pulsed with LPS-coated magnetic beads (A, C) or LPS-coated polystyrene beads (B, D) and chased as indicated. A. Purified phagosomes (*left*) or whole cell lysates (*right*) were analyzed by SDS-PAGE and immunoblotting for TLR4 and actin. Shown are the relevant bands for each blot. C. Quantification of band intensities for phagosomal TLR4 from three independent experiments, showing fold change relative to each shRNA treatment at time 0 and normalized to actin (mean ± SD). (B, D). Phagosomes were purified and analyzed by flow cytrometry using a FITC-conjugated anti-TLR4 antibody. B. Shown are histogram plots of a representative experiment with the percentages of gated phagosomes that were TLR4 positive indicated. Solid black lines, non-target (control) shRNA; solid blue lines, PI4KIIβ shRNA; solid red lines, PI4KIIα shRNA; dashed lines, non-transduced controls. D. Data from three independent experiments performed in duplicate are shown as fold change of percent of TLR4**^+^** phagosomes relative to each shRNA treatment at time 0 (mean ± SD). *p<0.05; **p<0.01; ***p<0.001.

### 6. Sorting adaptor TIRAP recruitment to phagosomes is severely impaired by the knockdown of PI4KIIα

Formation of the MyDdosome complex and downstream TLR4 signaling is favored by TLR4 binding to its sorting adaptor TIRAP (Barnett & Kagan, 2019; Gay et al, 2011). TIRAP contains a N-terminal lysine-rich polybasic motif that promiscuously binds to different phosphoinositide species, including PtdIns(4,5)P_2_ and PtdIns4P (Bonham et al, 2014). While PtdIns(4,5)P_2_ mostly localizes to the plasma membrane and to nascent phagosomes, PtdIns4P – which is present on late endocytic compartments –, was a strong candidate for TIRAP binding to maturing phagosomal membranes. To test whether TIRAP is recruited to phagosomes and whether recruitment requires PI4KIIα, we followed the kinetics of TIRAP-GFP recruitment to phagosomes by live cell imaging (Figure 7) and flow cytometry on isolated phagosomes (Figure 8) from TIRAP-GFP transduced DCs (**Fig. EV2C**). In cells pulsed with OVA-TxR beads, TIRAP-GFP was detected on the plasma membrane and on nascent phagosomes, consistent with its ability to bind PtdIns(4,5)P_2_ and similar to the localization of the PtdIns(4,5)P_2_-sensing probe GFP-PH-PLCδ (pleckstrin homology domain of phospholipase Cδ; **Figure EV5A**, **B**). Recruitment of TIRAP-GFP and GFP-PH-PLCδ to the plasma membrane was modestly affected by knockdown of PI4KIIβ but not PI4KIIα, whereas TIRAP-GFP (but not GFP-PH-PLCδ) was detected on many fewer phagosomes in PI4KIIα knockdown cells after the pulse (Figure 7A, B time 0 and **Figure EV5A**, **B**). Over time, TIRAP-GFP was increasingly recruited to phagosomes in control and PI4KIIβ knockdown cells, as quantified both by fluorescence microscopy and flow cytometry on isolated phagosomes (Figure 7B, Figures 8A, C and **Movie EV8**) and was detected on phagotubules at 120 min (Figure 7A and **Movie EV9**). However, TIRAP recruitment to phagosomes was severely reduced in PI4KIIα knockdowns between 30 and 120 min (on 90±10% of phagosomes in control and PI4KIIβ knockdown cells vs. 1±5% of phagosomes in PI4KIIα knockdown DCs; Figure 7B and **Movie EV10**), with no detectable phagosomal increase as measured by flow cytometry on isolated phagosomes (Figure 8A, C). Consistent with the role of AP-3 in PI4KIIα recruitment to phagosomes, TIRAP recruitment to phagosomes was also impaired in AP-3-deficient DCs (1±5% of phagosomes between 30 and 120 min; Figure 7B). In contrast to these observations, phagosomal recruitment of the p40-phox domain containing PX-TIR-GFP construct, which preferentially binds PtdIns3P – a lipid enriched on early endosomes and early phagosomes (Vieira et al, 2001) – was not affected by PI4KIIα knockdown or AP-3 deficiency (Figure 8B, D-F). PX-TIR-GFP was mainly detected on phagosomes between 30 and 60 min after the pulse (Figure 8B, D-F), at times when the PtdIns4P probe GFP-P4Mx2 and TIRAP-GFP were also detected in control and PI4KIIβ knockdown DCs (Figures 2 and 7). The observation that PI4KIIα knockdown or AP-3 knockout severely impairs TIRAP but not PX-TIR recruitment to phagosomes suggest that TIRAP preferentially binds PtdIns4P on DC phagosomes. Note that phagotubules are labeled by OVA-TxR but not by PX-TIR-GFP (Figure 8E), suggesting that phagotubules, which emanate from mature phagosomes, do not contain PtdIns3P, a lipid mainly present on early phagosomes. These results show that PI4KIIα is necessary and sufficient for TIRAP recruitment to phagosomes, and suggest that, in contrast to its recruitment to early endosomes, PtdIns3P is not sufficient to recruit TIRAP to early phagosomes in DCs.

**Figure 7.**
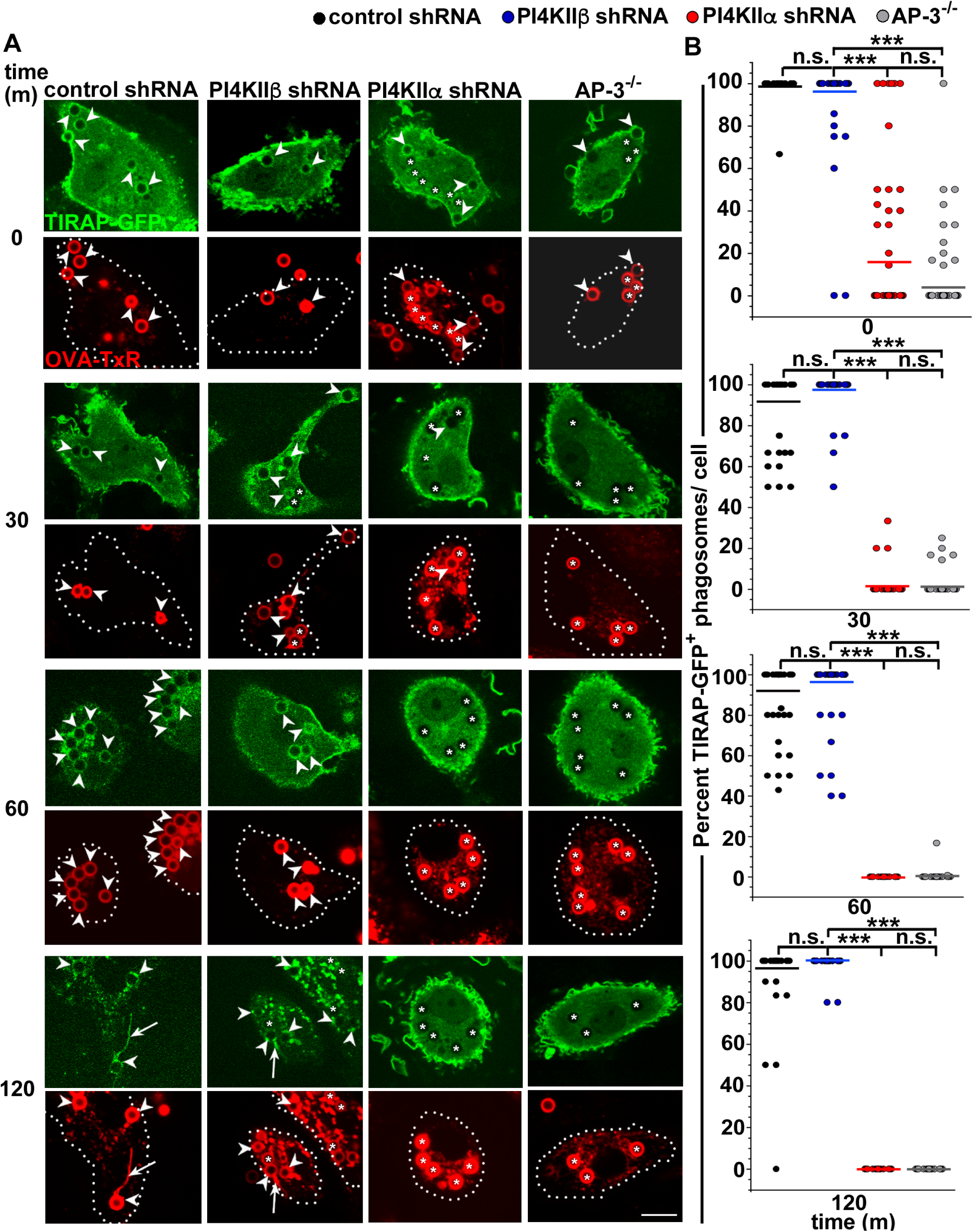
Sorting adaptor TIRAP recruitment to phagosomes is severely impaired by PI4KIIα knockdown. WT BMDCs were transduced with retroviruses encoding TIRAP-GFP and lentiviruses encoding non-target (control), PI4KIIβ or PI4KIIα shRNAs, and AP-3^-/-^ BMDCs were transduced only with retroviruses encoding TIRAP-GFP. DCs were pulsed with LPS/OVA-TxR-coated beads, chased as indicated and analyzed by live cell imaging. A. Representative images. Dotted white lines, cell outlines; arrowheads, TIRAP-GFP positive phagosomes; asterisks, TIRAP-GFP negative phagosomes; arrows, phagotubules. B. Data from three independent experiments, 20 cells per experiment, are presented as percent of TIRAP-GFP**^+^** phagosomes per cell. Black dots, non-target control shRNA; blue dots, PI4KIIβ shRNA; red dots, PI4KIIα shRNA; grey dots, AP-3^-/-^; solid color lines, means. Scale bar: 9 µm. ***p<0.001; n.s., not significant.

**Figure 8.**
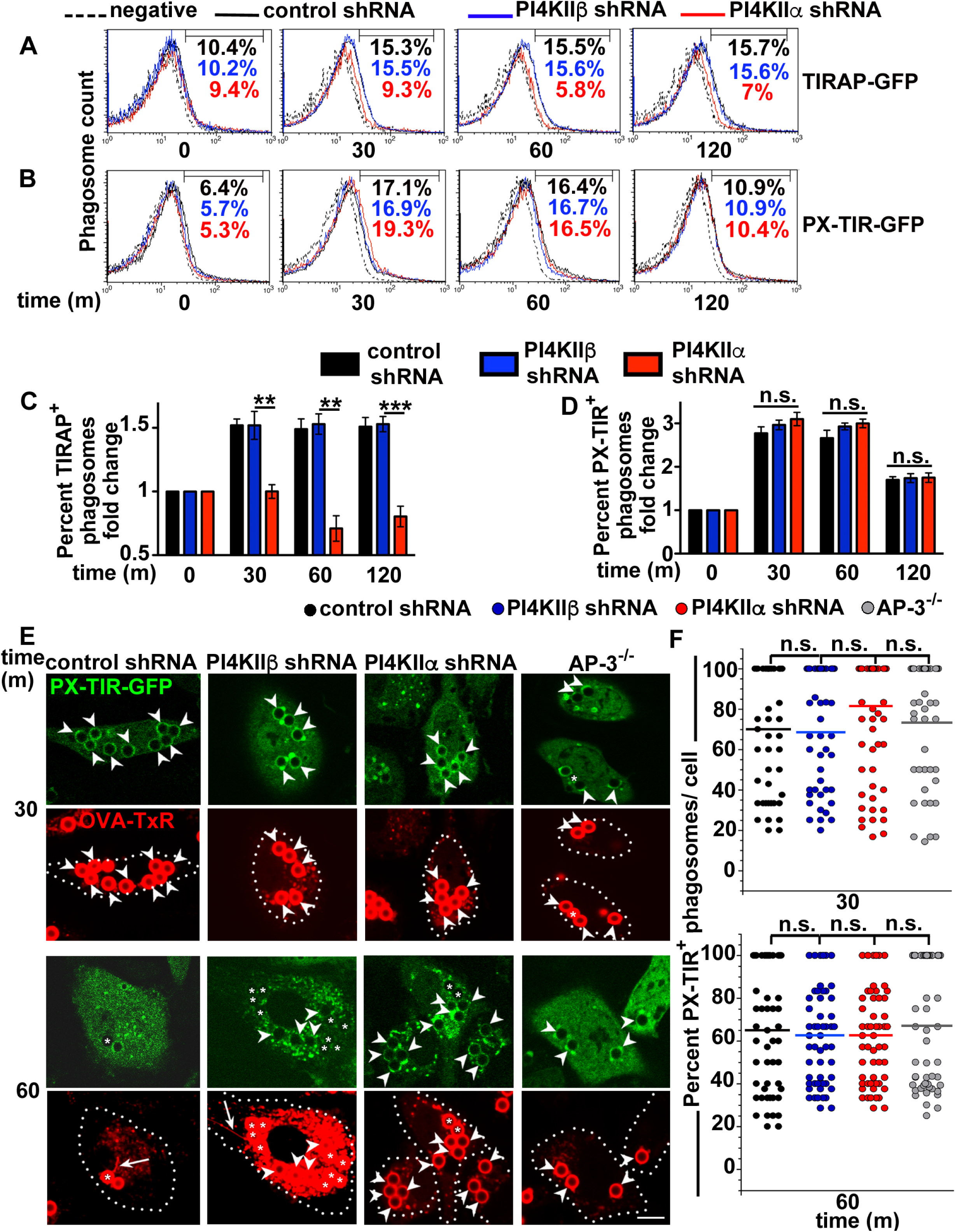
Phagosomal recruitment of the sorting adaptor TIRAP, but not of the PtdIns3P-binding PX-TIR, is severely impaired by PI4KIIα knockdown. WT BMDCs transduced with retroviruses encoding TIRAP-GFP or PX-TIR-GFP and lentiviruses encoding non-target (control), PI4KIIβ or PI4KIIα shRNAs, or AP-3^-/-^ BMDCs transduced only with retroviruses encoding TIRAP-GFP, were pulsed with LPS-coated polystyrene beads (A-D) or LPS/OVA-TxR-coated beads (E, F) and chased as indicated. (A-D). Phagosomes were purified and analyzed by flow cytometry. GFP-positive BMDCs were not previously sorted. (A, B). Shown are histogram plots of a representative experiment with the percentages of gated phagosomes that were GFP positive indicated. Solid black lines, non-target (control) shRNA; solid blue lines, PI4KIIβ shRNA; solid red lines, PI4KIIα shRNA; dashed lines, non-transduced controls. (C, D). Data from three independent experiments performed in duplicate are shown as fold change of percent of TIRAP-GFP**^+^** phagosomes (C), or PX-TIR-GFP positive phagosomes (D) relative to each shRNA treatment at time 0 (mean ± SD). (E, F). BMDCs expressing PX-TIR-GFP were analyzed by live cell imaging after 30 and 60m. E. Representative images. Dotted white lines, cell outlines; arrowheads, PX-TIR-GFP positive phagosomes; asterisks, PX-TIR-GFP negative phagosomes; arrows, phagotubules. Note that phagotubules are labeled by Tx-Red but not by PX-TIR-GFP. F. Data from three independent experiments, 20 cells per experiment, are presented as percent of PX-TIR-GFP positive phagosomes per cell. Black dots, non-target control shRNA; blue dots, PI4KIIβ shRNA; red dots, PI4KIIα shRNA; grey dots, AP-3^-/-^; solid color lines, means. Scale bar: 6 µm. **p<0.01; ***p<0.001; n.s., not significant.

## DISCUSSION

Intracellular trafficking pathways play a crucial role in preserving organelle identity. We previously showed that a key mediator of TLR4 recruitment from endosomes to phagosomes in DCs is the trafficking adaptor protein AP-3. However, loss of AP-3 expression did not completely abrogate TLR recruitment or signaling, suggesting that AP-3 might play a regulatory role in this process. We now provide evidence that AP-3 does not directly bind TLR4, suggesting that AP-3 regulates TLR4 localization to phagosomes in an indirect manner. We have now identified the lipid kinase PI4KIIα as an AP-3 cargo that is required to promote TLR4 signaling from phagosomes. PI4KIIα trafficking to phagosomes in DCs requires its kinase activity and binding to AP-3, consistent with the requirements for PI4KIIα delivery to lysosomes, lysosome related organelles, and synaptic vesicles in other cell types (Craige et al, 2008). PI4KIIα in turn generates PtdIns4P on phagosomes, and knockdown of PI4KIIα leads to reduced phagosomal PtdIns4P as measured by accumulation of the P4Mx2 probe. The phagosomal effects are specific for PI4KIIα, as knockdown of the genetically distinct PI4KIIβ – which does not bind AP-3 – does not impact phagosomal PtdIns4P or TLR4 signaling. Finally, we show that PI4KIIα activity is required to recruit the TLR sorting adaptor TIRAP to phagosomes, explaining PI4KIIα-dependent TLR4 accumulation on phagosomes. Together, the data suggest a model in which AP-3-dependent trafficking of PI4KIIα to phagosomes generates a pool of PtdIns4P that is necessary to recruit TIRAP and sustain TLR4 proinflammatory signaling and MHC-II presentation from phagosomes.

Our data show that PI4KIIα builds a PtdIns4P platform on the maturing phagosome that allows the binding of the TLR sorting adaptor TIRAP. TIRAP serves as a landmark for the assembly of the MyDdosome complex on phosphoinisotide-enriched domains, promoting the initiation of TLR signal transduction. TIRAP was shown to promiscuously bind to different phosphoinositides in macrophages and DCs, including PtdIns(4,5)P_2_ on the plasma membrane and PtdIns3P on early endosomes (Bonham et al, 2014). Here we show that on DC maturing phagosomes, TIRAP binding is highly dependent on PtdIns4P, and that knockdown of PI4KIIα, but not of PI4KIIβ, significantly reduces TIRAP recruitment to phagosomes. In contrast, TIRAP association with the plasma membrane and nascent phagosomes is PI4KIIα independent – similarly to PH-PLCδ, which detects PtdIns(4,5)P_2_ –, suggesting that TIRAP binding to the plasma membrane is not absolutely dependent on PtdIns4P and requires PtdIns(4,5)P_2._ Consistent with observations in macrophages, PtdIns3P (detected with PX-TIR) accumulates on early phagosomes in DCs, and its presence is not dependent on PI4KIIα or PI4KIIβ. However, TIRAP is not recruited to phagosomes at this stage in the absence of PI4KIIα, suggesting that TIRAP is also preferentially recruited to early phagosomes by PtdIns4P. This suggests that additional determinants might limit or promote phosphoinositide-dependent TIRAP recruitment to membranes, and highlights differences between the early endosomal and phagosomal systems in DCs. Moreover, these data support the importance of distinguishing TLR signaling from distinct subcellular locations to reflect differences in cargo characteristics or origins, in this case soluble versus particulate, consistent with our observations that AP-3 is required for TLR signaling induced by phagosomal but not endosomal cargoes (Mantegazza et al, 2012).

This distinction between the endosomal and phagosomal systems in DCs could also reflect variations between DC and other phagocyte endolysosomal systems. Based on differences in the progression of phosphoinositide content shown here for DCs and previously for macrophages (Jeschke et al, 2015; Levin et al, 2017), and on the acquisition of lysosomal proteins and low pH, the phagosomal maturation process is slowed in DCs compared to that in macrophages, supporting DC specialization in antigen presentation (Savina et al, 2006; Trombetta et al, 2003). Consistent with this, while PtdIns(4,5)P_2_ and PtdIns3P are present on nascent phagosomes and early phagosomes respectively in macrophages (Levin et al, 2017), both persist longer on phagosomes in DCs than in macrophages. By contrast, PtdIns4P is generated earlier in the phagosomal maturation process in DCs compared to macrophages. This earlier formation of PtdIns4P on DC phagosomes with the consequent recruitment of TIRAP and assembly of the MyDdosome licenses DC phagosomes for antigen presentation, an activity that is not a primary function in macrophages (Mantegazza et al, 2013; Savina & Amigorena, 2007).

Of note, even though knockdown of PI4KIIα abolished TIRAP binding to maturing phagosomes, TLR4 downstream responses were not completely abrogated. The persistence of TLR4 signaling in the absence of TIRAP phagosomal recruitment suggests residual signaling from plasma membrane engaged TLRs, which are not regulated by PI4KIIα or AP-3. Indeed, we show that PI4KIIα specifically contributes to the phagosomal PtdIns4P pool. In contrast, the plasma membrane pool of PtdIns4P – while somewhat reduced by knockdown of PI4KIIβ but not of PI4KIIα –, is primarily maintained by PI4KIIIα (Balla et al, 2005), supporting – together with the large pool of PtdIns(4,5)P_2_ – TLR4/Myddosome signaling from the plasma membrane in cells depleted of PI4KIIα. In addition, it is possible that TLR4 could signal on the phagosome independently of TIRAP. Indeed, TIRAP requirement for TLR signaling could be bypassed by high ligand concentrations that may not reflect physiological conditions (Bonham et al, 2014; Horng et al, 2002; Yamamoto et al, 2002).

In live cell imaging experiments, PI4KIIα, PtdIns4P and TIRAP were also detected on phagosomal tubules that we had previously shown to promote MHC-II presentation from phagosomes. Moreover, knockdown of PI4KIIα or loss of AP-3 expression abrogated phagosomal tubule formation. This could reflect reduced binding of the Rab7 adaptors RILP and FYCO1, which respectively modulate dynein and kinesin microtubule motor proteins required for tubule formation (Mrakovic et al, 2012). Indeed, Rab7 is present on PtdIns4P enriched compartments (Baba et al, 2019). It could also suggest that PtdIns4P are required for the recruitment of membrane curvature stabilizing proteins (Suetsugu et al, 2014). Expression of WT PI4KIIα, but not the kinase-inactive or AP-3 sorting mutants, rescued DC defects in phagosomal TLR signaling and MHC-II presentation. Together, these observations support the conclusion that AP-3-dependent recruitment of PI4KIIα to phagosomes generates a pool of PtdIns4P that is required for the formation of phagosomal tubules, TIRAP recruitment and concomitant enhancement of MHC-II presentation. Our data also suggest that impaired phagosomal PI4KIIα recruitment may at least in part explain defective anti-bacterial immune responses in AP-3 deficient mice and HPS2 patients.

In summary, our data indicate that AP-3-mediated recruitment of PI4KIIα early in the life cycle of the DC phagosome is a prerequisite for the binding of TIRAP and the promotion of TLR4 signaling, and is a key determinant of the fate of the phagosome as an autonomous signaling organelle. Further studies will be required to elucidate the signals that drive TIRAP and TLR4 together on the phagosome to allow MyDdosome formation.

## MATERIALS AND METHODS

### Mice

Mice were bred under pathogen-free conditions in the Department of Veterinary Resources at the Children’s Hospital of Philadelphia and were euthanized by carbon dioxide narcosis according to guidelines of the American Veterinary Medical Association Guidelines on Euthanasia. All animal studies were performed in compliance with the federal regulations set forth in the recommendations in the Public Health Service Policy on the Humane Care and Use of Laboratory Animals, the National Research Council’s Guide for the Care and Use of Laboratory Animals, the National Institutes of Health Office of Laboratory Animal Welfare, the American Veterinary Medical Association Guidelines on Euthanasia, and the guidelines of the Institutional Animal Care and Use Committees of Children’s Hospital of Philadelphia. All protocols used in this study were approved by the Institutional Animal Care and Use Committee at the Children’s Hospital of Philadelphia. C57BL/6 (CD45.1 or CD45.2) wild-type (WT) mice and OT-II TCR (vα2-vβ5)-transgenic mice were originally purchased from The Jackson Laboratories (Bar Harbor, ME). The HPS2 mouse model pearl (AP-3 deficient; B6Pin.C3-*Ap3b1^pe^*^/*pe*^) was previously described (Feng et al, 1999) and kindly provided by Susan Guttentag (Vanderbilt University, Nashville, TN). *Tlr4^-/-^* (Hoshino et al, 1999) mice were kindly provided by Susan Ross (University of Pennsylvania, with the permission of Shizuo Akira, Osaka University, Japan). All mice were bred and housed in the same room to minimize possible microbiota differences. Sex- and age-matched mice between 6 and 12 weeks of age were used in all experiments.

### Cell culture

Bone marrow cells were isolated and cultured for 7±9 d in RPMI-1640 medium (Gibco, ThermoFisher, Waltham, MA) supplemented with 10% low endotoxin defined FBS (Hyclone, Logan, UT), 2mM L-Gln, 50 µM 2-mercaptoethanol (InVitrogen) and 30% granulocyte-macrophage colony stimulating factor (GM-CSF)-containing conditioned medium from J558L cells (kindly provided by former Ralph Steinman laboratory and Maria Paula Longhi, Queen Mary University of London, UK) for differentiation to DCs as described (Mantegazza & Marks, 2015; Winzler et al, 1997). Maturation was induced by 18 hr treatment with LPS (0.1 µg/ml). CD4+ T cells were isolated from single cell suspensions of lymph nodes from OT-II transgenic mice, by positive selection with anti-CD4 microbeads (Miltenyi Biotec Inc., Auburn, CA).

### Reagents

Lipopolysaccharide (LPS) was purchased from Sigma-Aldrich (Saint Louis, MO)(*Escherichia coli* (0111:B4) and coated onto 3-micron polystyrene beads (Polysciences Inc., Warrington, PA) as previously described (Mantegazza & Marks, 2015). OT-II peptide OVA_323-339_ (ISQAVHAAHAEINEAGR) was from InvivoGen (San Diego, CA) and kindly provided by Paula Oliver (Children’s Hospital of Philadelphia, Philadelphia, PA), and Eα_52-68_ peptide (ASFEAQGALANIAVDKA) was from AnaSpec (Fremont, CA). EαGFP fusion protein was purified as described (Itano & Jenkins, 2003) from a construct kindly provided by Marion Pepper Pew (University of Washington, Seattle, WA) and Marc Jenkins (University of Minnesota, Minneapolis, MN). Ovalbumin (OVA, grade V) and bovine serum albumin (BSA, fraction V) were from Sigma-Aldrich (Saint Louis, MO). All oligonucleotides used were customized and ordered from Integrated DNA Technologies (Coralville, IA). All restriction enzymes were from New England Biolabs (Ipswich, MA). Polymerase chain reaction (PCR) was performed using GoTaq kit (Promega, Madison, WI) unless otherwise indicated.

### Antibodies

Rabbit polyclonal antibodies to PI4KIIα and PI4KIIβ (Guo et al, 2003) were kindly provided by Pietro De Camilli (Yale University, New Haven, CT) and Victor Faundez (Emory University, Atlanta, GA). The mouse monoclonal antibody Yae (clone eBioY-Ae) to the Eα_52-68_:I-A^b^ complex was purchased from eBioscience (San Diego, CA). Rabbit polyclonal antibodies to TLR4 (H-80, used for immunoblotting) was from Santa Cruz (Santa Cruz, CA). FITC-conjugated anti-mouse vβ5 5.1/ 5.2 TCR (MR9-4), PE-conjugated anti-mouse H-2K^b^ (AF6-88.5), FITC conjugated anti-mouse I-A^b^ (AF6-120.1), hamster anti-mouse CD3ε (145-2C11), PE-conjugated rat anti-mouse CD4 (GK1.5), APC-conjugated rat anti-mouse CD11b (M1/70), PE-conjugated hamster anti-mouse CD11c (HL3), hamster anti-mouse CD28 (37.51), PE-conjugated rat anti-mouse CD40 (3/23), APC-conjugated hamster anti-mouse CD69 (H1.2F3) and FITC-conjugated rat anti-mouse CD86 (GL1) were from BD Biosciences (San Diego, CA). FITC conjugated rat anti-mouse TLR4/CD284 mAb (MTS510) used in flow cytometric analyses was from Imgenex (San Diego, CA). Mouse monoclonal anti-β actin and anti-γ tubulin used for immunoblotting were from Sigma. Mouse monoclonal anti-GFP was from Roche (Indianapolis, IN). All secondary antibodies used for immunoblotting were Near-InfraRed fluorescent conjugates (Jackson Immunoresearch, West Grove, PA). ELISA anti-mouse IL-2 and IL-6 sets were from BD Biosciences.

### DNA constructs and shRNAs

Retroviral constructs MSCV2.2-TIRAP-GFP and MSCV2.2-PX-TIR-GFP were gifts from Jonathan Kagan (Harvard Medical School) (Bonham et al, 2014) (Addgene plasmids #52739 and #52737 respectively), MigR1 was kindly provided by Warren Pear (University of Pennsylvania) and human pEGFP-C1-PI4KIIα, pEGFP-C1-P4M-SidMx2 and pEGFP-C1-GFP-PH-PLCδ were previously described (Balla & Varnai, 2002; Hammond et al, 2014; Stauffer et al, 1998; Varnai & Balla, 1998). A novel and unique NotI restriction site was introduced upstream of the ApaI restriction site in the MigR1 vector using the forward primer 5’-GTTAA**GCGGCCGC**AA ATTAGGGCC-3’ (NotI restriction site in bold), and the reverse primer: 5’-CTAATTT**GCGGCCG C**TTAACGGCC-3’ (NotI restriction site in bold). The ApaI restriction site is no longer present in the MigR1-NotI vector. GFP-P4M-SidMx2 was then amplified using the forward primer: 5’-ACTGAT**GGATCC**ATGGTGAGCAAGGGCGAG-3’ (BamHI restriction site in bold), and reverse primer 5’-ATCTAT**GCGGCCGC**TTATTTTATCTTAATGG-3’ (NotI restriction site in bold), and subcloned into the BglII and NotI restriction sites of the MigR1-NotI vector. C-terminally GFP-tagged PI4KIIα and PH-PLCδ were subcloned into the XhoI and NotI sites of the MIGR1-NotI vector as follows. pEGFP-N1-PI4KIIα was digested with XhoI and NotI restriction enzymes and the insert directly ligated with XhoI/ NotI digested vector. GFP-PH-PLCδ was amplified using the forward primer 5’-TTCA**CTCGAG**CTCAAGCTTCGAATTCTGCAGTC-3’ (XhoI restriction site in bold), and the reverse primer: 5’ATCT**GCGGCCGC**TTACTGGATGTTGA GCTC-3 (NotI restriction site in bold). The product was digested with NotI and XhoI for subcloning. The AP-3 sorting mutant PI4KIIα(L61,62A)-GFP and the kinase-inactive mutant PI4KIIα(D308A)-GFP were previously described (Craige et al, 2008). The AP-3 sorting mutant was regenerated in MigR1-PI4KIIα-GFP using the Q5 site-directed mutagenesis kit (New England Biolabs) following manufacturer’s instructions. Primers used in the L61-62A mutagenesis reaction were 5’-GCGGCAGCCAGCGGCGGATCGGGCCCGGGGCGC-3’ and 5’-TCGCGGTCGTGGCCCGGC 3’. The kinase-inactive mutant was generated by a two-step PCR method using the Phusion High-fidelity kit (New England Biolabs). In the first step, PI4KIIα base pairs 1-936 and the sequence containing the rest of PI4KIIα (915-1437) and GFP were amplified using the following set of primers: Forward primer 5’ CCGCTCGAGATGGACGAGAC GAG-3’ and reverse primer 5’-GTCATTGCCGCGCGCAGTGTTG-3’ for the 1-936 region, and forward primer 5’-CAACACTGCGCGCGGCAATGAC-3’ and reverse primer 5’-ATAAGAATGCGGCCG CTTTACTTGTACAG-3’ for the 915-1437 region. The two PCR products contain a 22-base-pair overlap region containing the D308A mutagenized sequence (CAACACTGCGCGCGGCAAT GAC). The second PCR concatenated the two PCR products to amplify full-length PI4KIIα (D308A)-GFP using the forward primer 5’-CCGCTCGAGATGGACGAGACGAG-3’ and the reverse primer 5’-ATAAGAATGCGGCCGCTTTACTTGTACAG-3’. PI4KIIα(D308A)-GFP was then subcloned into the XhoI and NotI sites of the MiGR1-NotI vector.

pLKO.1-puromycin derived lentiviral vectors (Stewart et al, 2003) for small hairpin RNAs (shRNAs) against PI4KIIα, PI4KIIβ, sec22b and non-target shRNAs were obtained from the High-throughput Screening Core of the University of Pennsylvania. PI4KIIα #1 sense sequence: CCCAAGAATGAAGAGCCATAT; PI4KIIα #2 sense sequence: CCGTTCTCTCAGGAGATCAA A; PI4KIIβ sense sequence: GCTGTTTGTGAAAGATTACAA; sec22b sense sequence: CCCTA TTCCTTCATCGAGTTT; non-target sense sequence: GCGCG ATAGCGCTAATAATTT.

### Lentiviral and retroviral production, and transduction of dendritic cells

Recombinant retroviruses encoding GFP-tagged fusion proteins were produced by transfection of the packaging cell line PLAT-E (Morita et al, 2000) (a generous gift of Mitchell Weiss, St. Jude Children’s Research Hospital, Memphis, TN) using Lipofectamine 2000 (Invitrogen, Thermo-Fisher Scientific), and harvesting from cell supernatants 2 d later. 3 x 10^6^ BM cells were seeded on 6-well non-tissue culture treated plates per well for transduction 2 d after isolation, and transduced by spinoculation with 3 ml of transfected Plat-E cell supernatant in the presence of 8 µg/ml polybrene and 20 mM HEPES for 2h at 37°C. Retrovirus-containing media were then replaced with DC culture media. Cells were passaged 3 d after infection and were collected for experiments 3 d later.

Recombinant lentiviruses encoding shRNAs were produced by co-transfection of 293T cells (obtained from American Type Culture Collection, Mannassas, VA) with packaging vectors pDM2.G and pSPAx2 using calcium phosphate precipitation (Marks et al, 1995), and harvested from cell supernatants 2 d later. 3 x 10^6^ BM cells were seeded in 6-well non-tissue culture treated plates per well for transduction two d after isolation, and transduced by spinoculation with 3 ml of transfected 293T cell supernatant in the presence of 8 µg/ml polybrene and 20mMHEPES for 2h at 37°C (Savina et al, 2009). Lentivirus-containing media were then replaced with DC culture media. Puromycin (2 µg/ml) was added 3 d after infection, and cells were collected for experiments 3 d later.

For BM cell transduction with both retroviral constructs and lentiviral shRNAs, cells were first transduced with the indicated retroviral constructs, washed and subsequently transduced with the indicated lentiviral shRNAs. Lentivirus-containing media were then replaced with DC culture media. In order to select shRNA-expressing DCs, puromycin (2 µg/ml) was added 3 d after infection, and cells were collected for experiments 3 d later. Only lentiviral shRNAs are puromycin resistant.

### Yeast culture, transformation and two hybrid assays

The assay was performed essentially as described (Ohno et al, 1998). The *S. cerevisiae* strain HF7c (Clontech Laboratories, Takara Bio USA, Mountain View, CA), containing the reporters His3 and LacZ under the control of a GAL4 upstream activation sequence, was maintained on complete yeast extract/peptone/dextrose plates (Sitaram et al, 2012). The coding sequence for the cytoplasmic domain of TLR4 was amplified from TLR4 cDNA from Origene (Rockville, MD) using the forward primer: 5’-GAGCTCGAATTCGCCGCCACC AAGTATGGTAGAGGTG-3’, and the reverse primer: 5’-AGCTCTGTCGACTCAGATAGATGTTGCTTCC-3’, and then digested and subcloned into the EcoRI/ SalI sites of the pGBT9 plasmid for fusion to the GAL4 DNA-binding domain. pGBT9/TGN38 plasmid, containing the cytoplasmic domain of human TGN38 and pGADT7 plasmids containing the activation domain of GAL4 fused to the µ1, µ2 or µ3 subunits of AP-1, AP-2 or AP-3, respectively, were gifts from Juan Bonifacino (National Institutes of Health, Bethesda, MD) (Ohno et al, 1998). Cotransformation with pGBT9 and pGADT7 plasmids was performed by a modification of the lithium acetate procedure as described in the Yeast Protocols Handbook from Clontech. Plasmids expressing GAL4-activation domains and GAL4-DNA-binding domains contain the Leu2 and Trp1 selection markers, respectively, allowing the transformed HF7c yeast cells to grow on plates that do not contain leucine and tryptophan. If the fusion proteins interact, yeast cells will express the His3 reporter gene, allowing them to grow on plates that do not contain histidine. HF7c transformants were selected by spreading on plates lacking leucine and tryptophan. For colony growth assays, HF7c transformants were pooled and spotted once or in fivefold serial dilutions on plates lacking leucine, tryptophan, and histidine and allowed to grow at 30°C for 3–5 d.

### Phagosome purification and protein recruitment

Phagosomes were isolated essentially as described (Savina et al, 2010). Briefly, BMDCs were incubated for 15 min with LPS-coated latex beads or 3 µm magnetic beads (Dynabeads M-280 streptavidin, Invitrogen, NY) and then chased. Magnetic and nonmagnetic phagosomes were purified after different chase times by means of a magnet or differential centrifugation, respectively, as described (Guermonprez et al, 2003; Mantegazza et al, 2008). Purified LPS-bead phagosomes were fixed and stained with antibodies to TLR-4 or negative controls, or left unstained in the case of the GFP-expressing DCs and analyzed concurrently by flow cytometry, gating on the bead population (Savina et al, 2010) and by normalizing to the total percentage of GFP-positive cells in the case of the transduced DCs. Flow cytometry was performed using a FACSCalibur and CellQuest software (BD Biosciences, San Diego, CA). Protein extracts from purified magnetic phagosomes were analyzed by immunoblotting.

### Immunoblotting

Immunoblotting was performed essentially as described. Briefly, Laemmli sample buffer with 2-mercaptoethanol was added to protein lysates from phagosome purification and whole cell lysates. Samples were then fractionated by SDS-PAGE on 10% polyacrylamide gels, transferred to PVDF membranes (Immobilon-FL, Millipore) and analyzed using Alexa Fluor 680 or 790-conjugated secondary antibodies (Jackson ImmunoResearch) and Odyssey imaging system (LI-COR, Lincoln, NE). Densitometric analyses of band intensity was performed using NIH Image J software, normalizing to control protein levels.

### Cytokine secretion after TLR4 stimulation

BMDCs were incubated with LPS-coated beads for 3 hr as described (Mantegazza et al, 2012). IL-6 concentration in culture supernatants was measured by ELISA (BD Biosciences).

### Live cell imaging

BMDCs expressing retroviral and/or lentiviral constructs were seeded on poly-L-lysine–coated glass-bottom 35-mm culture dishes (MatTek, Ashland, MA) on day 7 of culture. On day 8, DCs were pulsed for 30 min with TxR-conjugated OVA (Invitrogen, ThermoFisher scientific) and LPS (100 µg/mL) covalently coupled to 3-µm amino polystyrene beads (Polysciences Inc., Warrington, PA) as described previously (Mantegazza & Marks, 2015). DCs were then washed with RPMI, chased for 0– 2.5 h, and visualized by spinning-disk confocal microscopy using an Olympus inverted microscope equipped with an environmental chamber at 37°C and 5% CO2 at the University of Pennsylvania’s Confocal Microscopy core or a BioVision (Milpitas, CA) and DMi8 Leica (Morrisville, PA) spinning-disk system using an Andor 888 cooled EMCCD camera equipped with temperature and CO_2_ control units, and associated VisiView software (Visitron Systems GmbH, Puchheim, Germany) for image and video capture. Time-lapse microscopy was performed by capturing image streams over 1–5 min at 1 frame/s and analyzed using ImageJ (National Institutes of Health, Bethesda, MD). Protein recruitment to phagosomes was visualized with the ImageJ plugin 3D viewer and quantified using Analyze/Plot profile and Analyze/3D surface plot as detailed in the ImageJ tutorial (https://imagej.nih.gov/ij/docs/menus/analyze.html).

### Antigen presentation assays

DCs were exposed to OVA, OVA:BSA-coated 3 µm latex beads (Polysciences, Warrington, PA), or OVA-specific MHC-II peptides for 15–30 min at 37°C, then washed in PBS and chased in complete medium at 37°C. DCs were then fixed with 0.005% glutaraldehyde in PBS for 1 min., washed with 0.2M glycine in PBS, and co-cultured with CD4+ OT-II T cells that had been prestimulated with anti-CD3 and anti-CD8 antibodies (Fitch et al, 2006). T cell activation was monitored 18 h later as CD69 expression by flow cytometry (FACSCalibur, BD Biosciences, San Diego, CA) and IL-2 secretion in coculture supernatants by ELISA (BD Biosciences). For presentation of Eα_52-68_ peptide on I-A^b^, BMDCs were incubated with 0.5 mg soluble EαGFP, Eα_52-68_ peptide or EαGFP-coated (1 mg/ml) beads for 30 min, washed with PBS, and chased. Cells were then fixed with 3% paraformaldehyde in PBS, stained with biotinylated YAe and APC-streptavidin (Invitrogen), and analyzed by flow cytometry. YAe labeling was quantified on CD11c+ cells that had taken up one bead as gated on a forward scatter versus side scatter plot (Lee et al, 2010).

### Statistical analyses

Statistical analyses and data plots were performed using Microsoft Excel (Redmond, WA) and GraphPad Prism software (San Diego, CA). Significance for experimental samples relative to untreated or non-target shRNA-treated WT control (unless otherwise stated) was determined using the unpaired Student’s t test and ANOVA. Mean ± SEM values are indicated in the main text. Error bars in figures represent mean ± SD.

## ACNOWLEDGMENTS

We thank Pietro De Camilli, Victor Faundez, Juan Bonifacino, Warren Pear, Marion Pepper Pew, Mark Jenkins, Paula Oliver, Susan Ross, Maria Paula Longhi and the former Ralph Steinman laboratory for the generous gifts of reagents, Anand Sitaram for experimental assistance, Andrea Stout and the Microscopy core and David Schultz and the High-throughput Screening core at the University of Pennsylvania for expert technical assistance and the Flow Cytometry core at the Children’s Hospital of Philadelphia. This work was supported by National Institutes of Health grants R01 AI137173 (to C.L.-H. and A.R.M.) and R01 HL121323 (to J.B.-K., Y.Z. and M.S.M.), Canadian Institutes of Health Research grant FDN-143202 (to R.L.-K. and S.G.), and the intramural research program of the Eunice Kennedy Shriver National Institute of Child Health and Human Development of the NIH (to T.B.).

## AUTHOR CONTRIBUTIONS

S.G., M.S.M. and A.R.M. conceived the project. T.B. provided phosphoinositide-detection probes and interpreted data. C.L.-H., M.S.M. and A.R.M. designed the experiments and analyzed the data. C.L.-H., R.L.-K., J.B.-K., Y.Z. and A.R.M. performed experimental work. A.R.M. drafted the manuscript. All authors reviewed and edited the manuscript.

## EXPANDED VIEW FIGURE AND MOVIE LEGENDS

**Figure EV1. AP-3 µ3 subunit does not bind the TLR4 cytoplasmic domain.** A yeast two-hybrid assay using Gal4 binding domains fused to the cytoplasmic domains of TGN38 or TLR4 or to nothing (empty vector) co-expressed with Gal4 activation domains fused to the µ1, µ2 or µ3 subunit of AP-1, AP-2 and AP-3, respectively. A protein–protein interaction leads to expression of *HIS3*, allowing growth on His-deficient medium. Top, triplicate analyses of yeast grown on medium lacking leucine and tryptophan but containing histidine (+His). Bottom, triplicate analyses of identical colonies grown on medium lacking leucine, tryptophan, and histidine (-His). Data are representative of 3 independent experiments.

**Figure EV2. Retroviral transduction efficiency is similar for all cell types and shRNA-expressing cells.** WT or AP-3^-/-^ BMDCs were non-transduced or transduced with retroviruses encoding PI4KIIα-GFP (A, B) and pulsed with LPS-coated polystyrene beads (B). (A). Shown are representative dot plots with the percentages of gated GFP-posiive DCs indicated. B. Shown are representative dot plots of beads (*left*) or purified phagosomes from WT DCs (*right*) indicating gated region (R1) based on side scatter (SSC) and forward scatter (FSC). C. BMDCs were non-transduced or transduced with retroviruses encoding TIRAP-GFP and lentiviruses encoding non-target (control), PI4KIIβ or PI4KIIα shRNAs. Shown are representative dot plots with the percentages of gated DCs that were GFP positive indicated.

**Figure EV3. Lentiviral transduction does not significantly impair DC differentiation, maturation or phagocytic capacity.** WT or AP-3^-/-^ BMDCs that were non-transduced (-in C) or WT BMDCs transduced with lentiviruses encoding non-target (control), PI4KIIβ, Sec22b or either of two PI4KIIα shRNAs were untreated (A, B) or pulsed with LPS-coated polystyrene beads for 20 min (C), or treated with soluble LPS for 18 h (D). A. Whole cell lysates were analyzed by SDS-PAGE and immunoblotting for PI4KIIα, PI4KIIβ and tubulin. Shown are the relevant bands for each blot. Note the non-specific band that migrates slightly faster than PI4KIIβ and that is recognized by the anti-PI4KIIβ antibody. B. Representative dot plots with the percentages of CD11b**^+^**/CD11c**^+^** BMDCs indicated. C. Representative dot plots indicating the percentages of DCs that have phagocytosed beads based on SSC and FSC. Cells not exposed to beads (DCs - beads) are shown for comparison. D. Representative histograms with the percentages of CD40**^+^** or MHC-II**^+^** DCs as markers of DC maturation. Blue solid lines, untreated DCs; red solid lines, LPS-treated DCs; black solid lines, unstained controls.

**Figure EV4. Effect of lentiviral transduction on MHC-II expression after phagocytosis.** WT BMDCs that were non-transduced or transduced with lentiviruses encoding non-target (control), PI4KIIβ, Sec22b or PI4KIIα shRNAs were pulsed with EαGFP-coated beads for 6 h, and surface levels of MHC-II and CD11c were analyzed by flow cytometry. Shown are representative dot plots with the percentages of CD11c**^+^**/MHC-II**^+^** DCs and the MFI values for MHC-II indicated.

**Figure EV5. PH-PLCδ-GFP binding to PtdIns(4,5)P_2_ at the plasma membrane and nascent phagosomes is not affected by PI4KIIα or PI4KIIβ knockdown**. WT BMDCs transduced with retroviruses encoding PH-PLCδ-GFP and lentiviruses encoding non-target (control), PI4KIIβ or PI4KIIα shRNAs were pulsed with LPS/OVA-TxR-coated beads and immediately analyzed by live cell imaging. A. Representative images. Dotted white lines, cell outlines; arrowheads, PH-PLCδ-GFP-positive phagosomes; asterisks, PH-PLCδ-GFP-negative phagosomes. B. Data from three independent experiments, 20 cells per experiment, are presented as percent of PH-PLCδ-GFP positive phagosomes per cell. Black dots, non-target control shRNA; blue dots, PI4KIIβ shRNA; red dots, PI4KIIα shRNA; solid color lines, means; n.s., not significant. Scale bar: 6 µm.

**Movie EV1**. **PI4KIIα is recruited to phagosomes in WT DCs.** This 3D animation of a series of z-stack confocal microscopy images shows the localization of PI4KIIα-GFP (*green*) to phagosomal membranes after pulsing WT DCs with LPS/OVA-TxR beads (*not colored*) and chasing for 30 min. Purple arrows point to phagosomes.

**Movie EV2**. **Reduced recruitment of PI4KIIα to phagosomes in AP-3^-/-^ DCs.** This 3D animation of a series of z-stack confocal microscopy images shows the reduced localization of PI4KIIα-GFP (*green*) to phagosomal membranes after pulsing AP-3^-/-^ DCs with LPS/OVA-TxR beads (*not colored*) and chasing for 30 min. Purple arrows point to phagosomes.

**Movies EV3 and EV4**. **PI4KIIα is recruited to phagosomal tubules in WT DCs**. These movies show tubules emerging from maturing phagosomes after pulsing WT DCs with LPS/OVA-TxR beads and chasing for 120 min. Note that phagotubules are positive for both PI4KIIα-GFP *(green)* and OVA-TxR (*red*).

**Movie EV5**. **Formation of phagosomal tubules is not affected by knockdown of PI4KIIβ.** This movie shows tubules emerging from maturing phagosomes after pulsing PI4KIIβ knockdown DCs with LPS/OVA-TxR beads (*red*) and chasing for 120 min.

**Movie EV6**. **Knockdown of PI4KIIα severely impairs phagosomal tubule formation.** This movie shows the absence of phagosomal tubules after pulsing PI4KIIα knockdown DCs with LPS/OVA-TxR beads (*red*) and chasing for 120 min.

**Movie EV7**. **AP-3 deficiency severely impairs phagosomal tubule formation.** This movie shows the absence of phagosomal tubules after pulsing AP-3^-/-^ DCs with LPS/OVA-TxR beads (*red*) and chasing for 120 min.

**Movie EV8**. **Sorting adaptor TIRAP is recruited to phagosomes in WT DCs.** This 3D animation of a series of z-stack confocal microscopy images shows the localization of TIRAP-GFP (*green*) to phagosomal membranes after pulsing WT DCs with LPS/OVA-TxR beads *(not colored)*. Purple arrows point to phagosomes.

**Movie EV9**. **Sorting adaptor TIRAP is recruited to phagosomal tubules in WT DCs.** This 3D animation of a series of z-stack confocal microscopy images shows the localization of TIRAP-GFP (*green*) to phagosomal membranes and phagotubules after pulsing WT DCs with LPS/OVA-TxR beads *(not colored)* and chasing for 120 min. Green arrows point to phagosomes.

**Movie EV10**. **Reduced recruitment of sorting adaptor TIRAP to phagosomes in PI4KIIα knockdown DCs.** This 3D animation of a series of z-stack confocal microscopy images shows the reduced localization of TIRAP-GFP (*green*) to phagosomal membranes, and the absence of phagotubules, after pulsing PI4KIIα knockdown DCs with LPS/OVA-TxR beads (*not colored*) and chasing for 120 min. Purple arrows point to phagosomes.

